# Introduction to Single-cell Physiologically-Based Pharmacokinetic (scPBPK) Models

**DOI:** 10.64898/2026.03.09.710595

**Authors:** Anshul Saini, James M. Gallo

**Affiliations:** Department of Pharmaceutical Sciences, School of Pharmacy and Pharmaceutical Sciences, Buffalo, NY; Department of Systems Biology, Columbia University, New York, NY

## Abstract

The current investigation introduces single-cell physiologically-based pharmacokinetic (scPBPK) models to gain insight into drug disposition at the cellular scale. The transition from standard PBPK (sPBPK) models to scPBPK models required depiction of expression-dependent (ED) processes, such as drug metabolism or membrane transport. ED processes utilize weighting functions – a defined or data-driven distribution –that yield heterogeneity in individual cell kinetics. Two scPBPK model examples are provided, one involving a drug (AZD1775) subject to 3 ED blood-brain barrier transport processes, and another drug (midazolam) with a single ED process of metabolism by hepatocytes. For both examples, the weighting function for each ED process was defined by a negative binomial distribution that is often used in scRNAseq analytics. The AZD1775 model simulations indicated a large degree of single cell drug concentration heterogeneity, whereas those for midazolam did not, due to high membrane transport relative to metabolism. scPBPK models offer a means to probe cellular pharmacokinetics compatible with modern omic technologies and may be extended to pharmacodynamic models.

**Teaser:** The modeling framework to predict drug concentrations in single cells is presented.

## INTRODUCTION

Physiologically-based PK (PBPK) models – herein referred to as standard PBPK (sPBPK) models - have existed for decades (1) and have become a cornerstone of mechanistic pharmacology and drug development (2, 3). sPBPK models enable drug kinetics to be examined in target organs either as lumped compartments or subdivided into homogeneous vascular, interstitial and intracellular compartments. Given the mechanistic underpinnings of sPBPK models, they provide a rational basis to design therapeutic drug regimens and characterize drug-drug interactions (4,5). Nonetheless, given the emphasis on sPBPK models, there has yet to be a means to develop single-cell PBPK (scPBPK) models.

sPBPK modeling approaches have been relatively dormant compared to advances in multiomic technologies that yield large transcriptomic, epigenetic and proteomic datasets to accelerate an understanding of the biology underlying diseases (6-8). Analyses of multiomic datasets enable construction of networks consisting of gene and protein nodes connected in biologically meaningful ways that provide insights into disease origins and therapeutic treatments, in addition to a rich set of associated hypotheses. Network biology has spawned the computational biology and quantitative pharmacology fields, often referred to as quantitative systems pharmacology (QSP) (9 - 11). By accounting for drug action or pharmacodynamics (PDs) QSP models are intimately linked to sPBPK models. Therefore, the delineation of scPBPK models is viewed as a necessary first step to decipher single cell pharmacokinetic/pharmacodynamic (PK/PD) characteristics.

## Results

### Fundamental Approach

A simple and modified generic PBPK model (Fig. 1) illustrates the essential composition of a scPBPK model. First, it can be recognized that the structure of the model is analogous to a sPBPK model with arterial blood or plasma flow entering a tissue and venous flow exiting the tissue. Ordinary differential equations (ODEs) are used to describe drug dynamics in each tissue while conserving blood flow and mass wherein each tissue has either a 1-, 2- or 3-subcompartment structure. The standard 1-compartment structure – known as a blood flow-limited model - lumps the vascular, interstitial fluid and intracellular subcompartments into one homogeneous compartment. Second, the liver (Fig. 1) represented as a 3-subcompartment structure contains an expression-dependent (ED) process, in this case, drug metabolism, in the intracellular space. To achieve single cell drug dynamics, both single cells and bulk cells are depicted in the intracellular compartment. By accounting for both single cell and bulk cells, all dynamic processes, membrane transport or flux and metabolism are conserved in the organ.

**Fig. 1.**
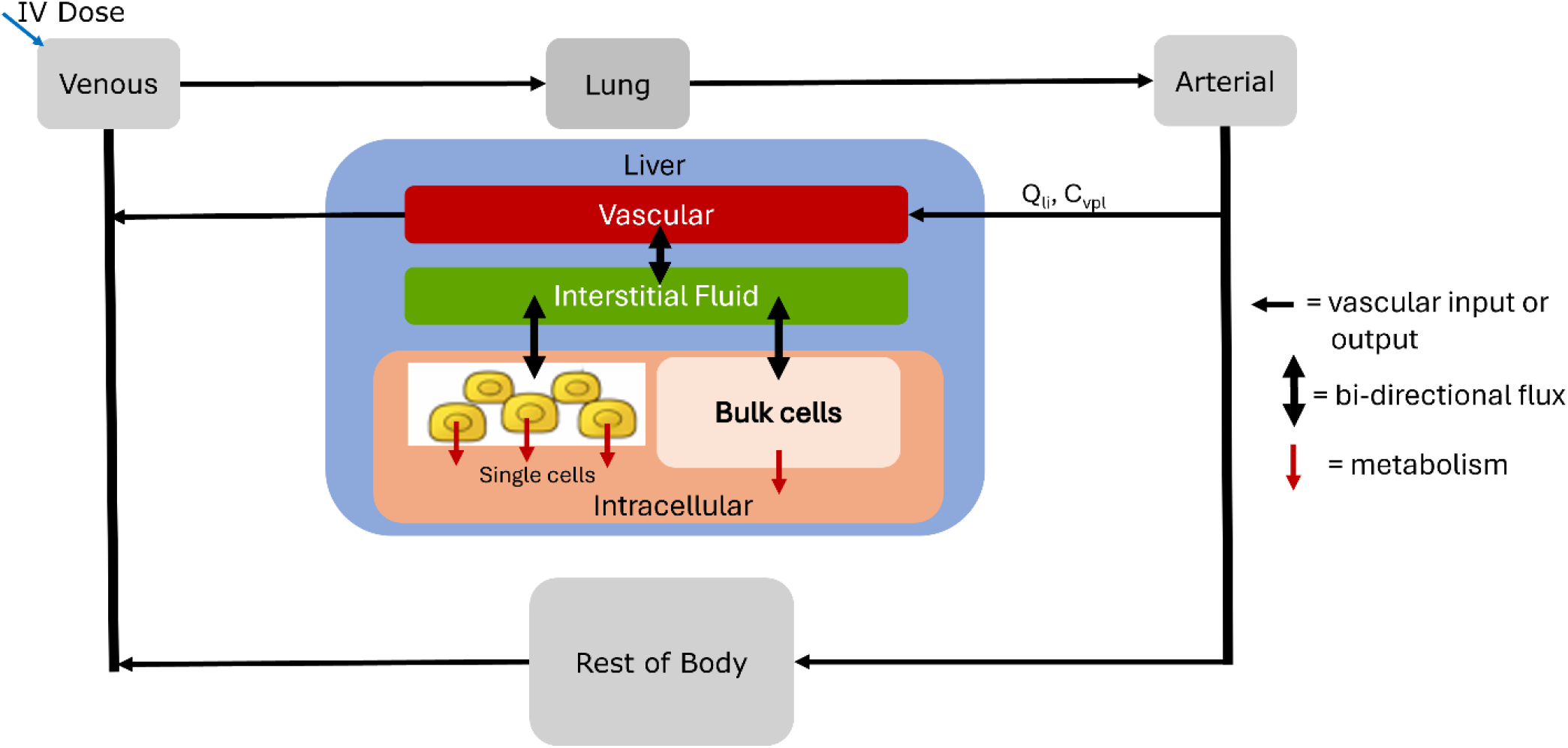
Illustration of a hypothetical scPBPK model. The liver depicted as a 3-subcompartment structure contains the expression-dependent (ED) process – metabolism – in the intracellular subcompartment. The intracellular compartment is divided into single cells that contain the ED process, and bulk cells.

Finally, a key aspect of casting scPBPK models is to identify expression-dependent (ED) processes, such as membrane transport or metabolism. Each ED process will incorporate a weighting function into the equation that accounts for the cellular heterogeneity of the unique parameter that is dependent on the expression of the protein or gene transcript. In the hypothetical scPBPK model (Fig. 1), drug metabolism is assumed to be ED and in a Michaelis-Menten metabolic reaction, the maximum metabolic capacity or *V*_*max*_, is considered to exhibit cellular heterogeneity due to variable protein abundances of the metabolic enzymes at the single cell level.

The ODEs for this hypothetical model are provided here:

Venous blood compartment (vbl):

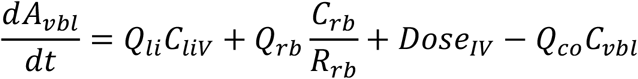

Lung compartment (lu):

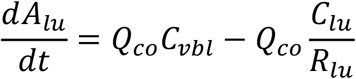

Arterial blood compartment (abl):

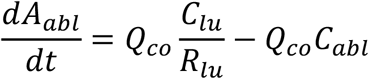

Rest of Body (rb):

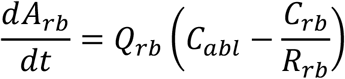

Liver compartment (li):

Vascular (V) subcompartment:

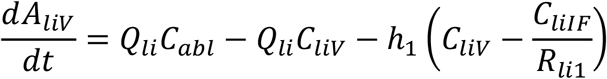

Interstitial Fluid (IF) compartment:

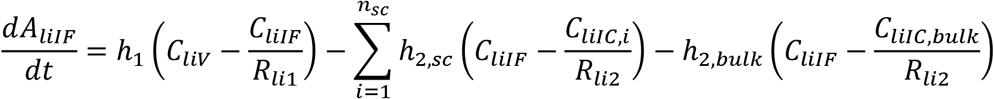

Intracellular (IC) compartment:

Single cell:

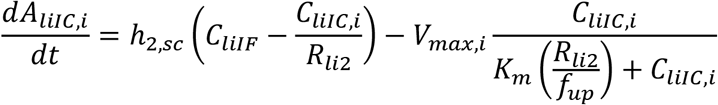

Bulk cells:

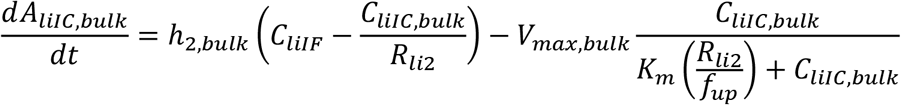

Drug concentrations are defined as:

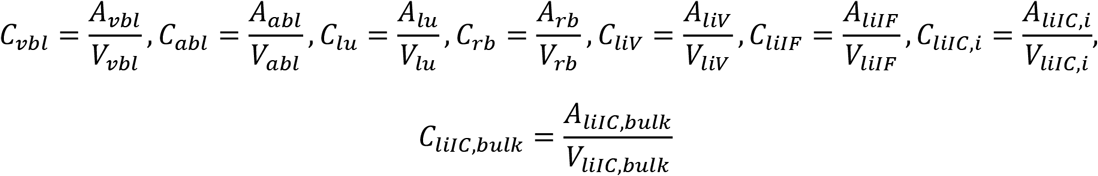

A number of relationships are defined for the liver ODEs:

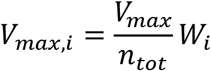

and

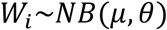

Each single cell has a unique *V*_*max,i*_ that is defined by the total *V*_*max*_ divided by the total number of hepatocytes, *n*_*tot*_, multiplied by, *W*_*i*_, a weighting function. The weighting function characterizes the expression level of the particular enzyme involved in the Michaelis-Menten reaction. For this hypothetical example, the weighting function is a negative binomial (NB) distribution with parameters, *𝜇*, the mean, and *𝜃*, the dispersion that is often used to analyze scRNAseq data (12, 13).

Also, in the single cell ODE:

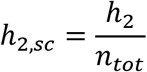

Each cell’s mass transport by diffusion, *h*_2,*sc*_, has to account for its particular contribution to mass transport, and therefore, the total mass transport, *h*_2_, is divided by the total number of cells, *n*_*tot*_.

Finally, in the bulk cell compartment ODE, two parameters require definition:

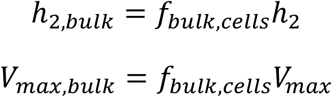

and

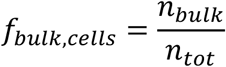

A conservation of kinetic processes – transport and clearance – are maintained when subcellular compartments are divided into single and cell bulk configurations. For instance, the total mass transport, *h*_2_, is equal to the sum of all the single cells and bulk processes:

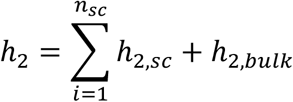

which can be rewritten as

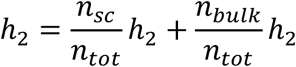

that is

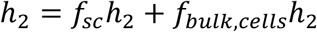

or *h*_2_ = *h*_2_, since *f*_*sc*_ + *f*_*bulk,cells*_ = 1.

This same conservation across single cells and bulk cells also applies to ED processes, wherein the weighting function, *W*_*i*_, yields the unique value of *V*_*max,i*_ for each cell that when summed over the number of single cells and bulk cells provides the total contribution of all single cells to the metabolic capacity. Thus, for *V*_*max*_,

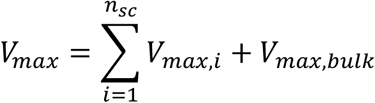

Table 1 below defines the terms in the ODEs and their units.

**Table 1.**
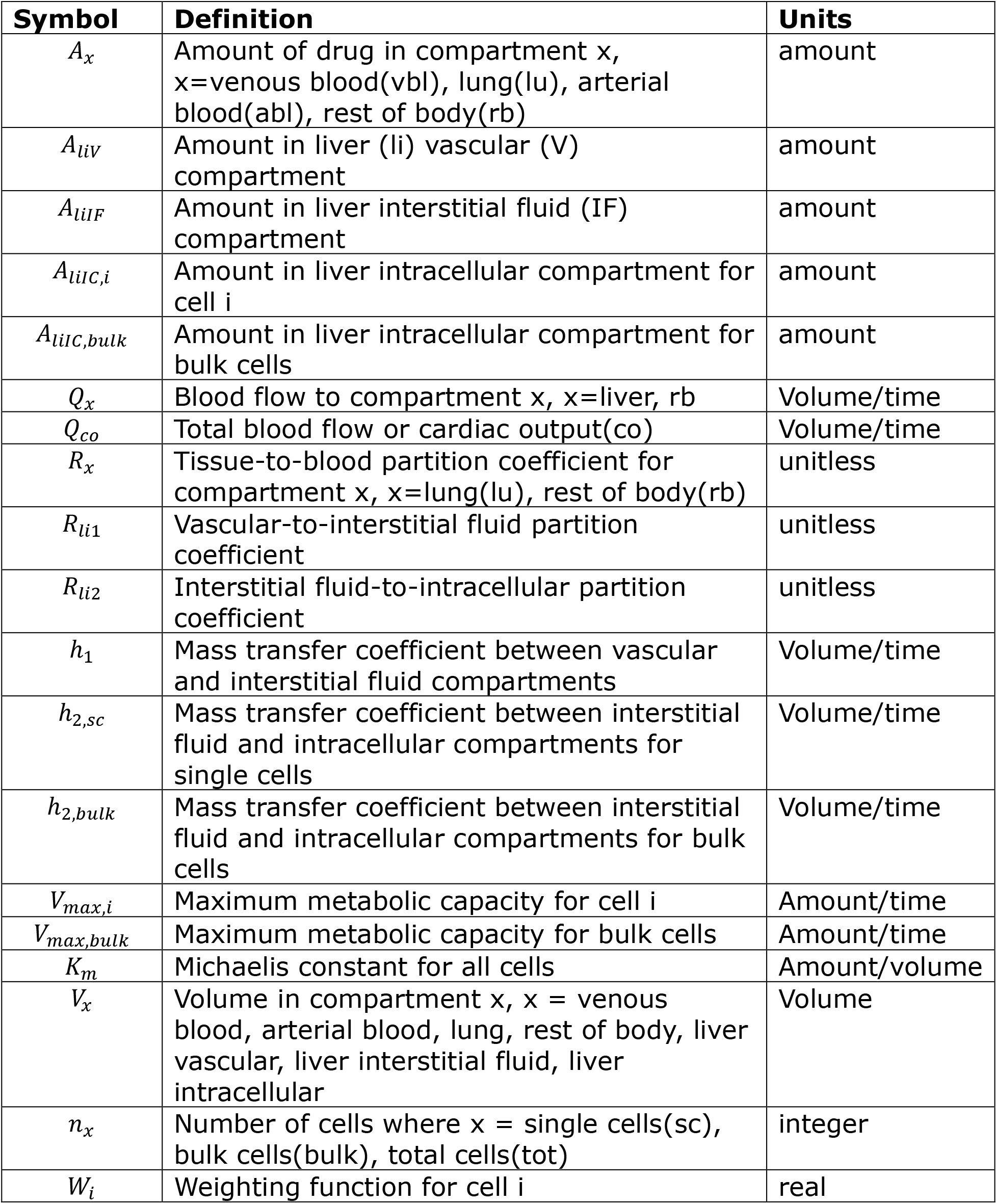
Definition of parameters in hypothetical scPBPK model.

There are two assumptions that warrant further consideration. First, it can be seen in the above equations that the total number of cells in the intracellular compartment, *n*_*tot*_, is required as are the number of single cells, *n*_*sc*_, and their difference defines the number of bulk cells, *n*_*bulk*_. There are sources to estimate cell numbers in tissues, such as for the number of hepatocytes in human livers (14), albeit the cell numbers can vary between individuals. Nonetheless, this is a parameter that can be set for each set of simulations. The single cell number, *n*_*sc*_, must be set and would be based on the number of cells that reflect the heterogeneity of each ED process as well as single cell data measurements of the implicated gene transcripts. In the simulations conducted for AZD1775 and midazolam, we selected *n*_*sc*_=10000 cells comprised of 4 clusters of 2500 cells each, which is consistent with cell numbers analyzed in scRNAseq experiments.

The weighting function, *W*_*i*_, has to be defined for each simulation. It should reflect the cellular heterogeneity of the designated ED process and can be determined from scRNAseq studies or as we did assume to conform to known statistical distributions, such as the negative binomial distribution, which can also clarify interpretation of transcriptomic data (12,13).

The unique steps to develop a scPBPK model are summarized:

1. Identify organs with ED processes.
2. Write rate equations for ED processes and define weighting function for each.
3. Define ODEs for system and utilize 3-subcompartment structure for organs containing ED processes.
4. Define cell numbers for those cells involved in ED processes.

### Standard PBPK Model for AZD1775

A standard PBPK (sPBPK) model for AZD1775 was used as the base model prior to converting it to a scPBPK model. The sPBPK model was developed from a previous PBPK model (15) for AZD1775, a WEE1 kinase inhibitor, that focused on its blood-brain barrier (BBB) permeability in brain tumor patients. AZD1775 (adavosertib) continues to be evaluated in a variety of cancers in combination therapy (16, 17).

The whole-body sPBPK model of AZD1775 is shown in Fig. S1, and the specific brain structure used by Li *et al* (15) is presented in Fig. 2. Here the brain is represented as a 2-subcompartment model, separated by the BBB into vascular and extravascular brain compartments. There are three ED processes and 1 expression independent process (PSB) at the BBB. Two of the ED processes are drug efflux pumps - P-glycoprotein (Pgp) and ABCG2 – known to restrict drug entry into the brain (18,19). The third ED process is an unspecified uptake transporter yet inferred to be OATP1B1. The bidirectional arrow labeled PSB represents passive diffusion cast as a permeability-surface area product. The other organs in the AZD1775 PBPK model were represented as blood flow-limited compartments as used by Li *et al* (15) and shown in Fig. S1.

**Fig. 2.**
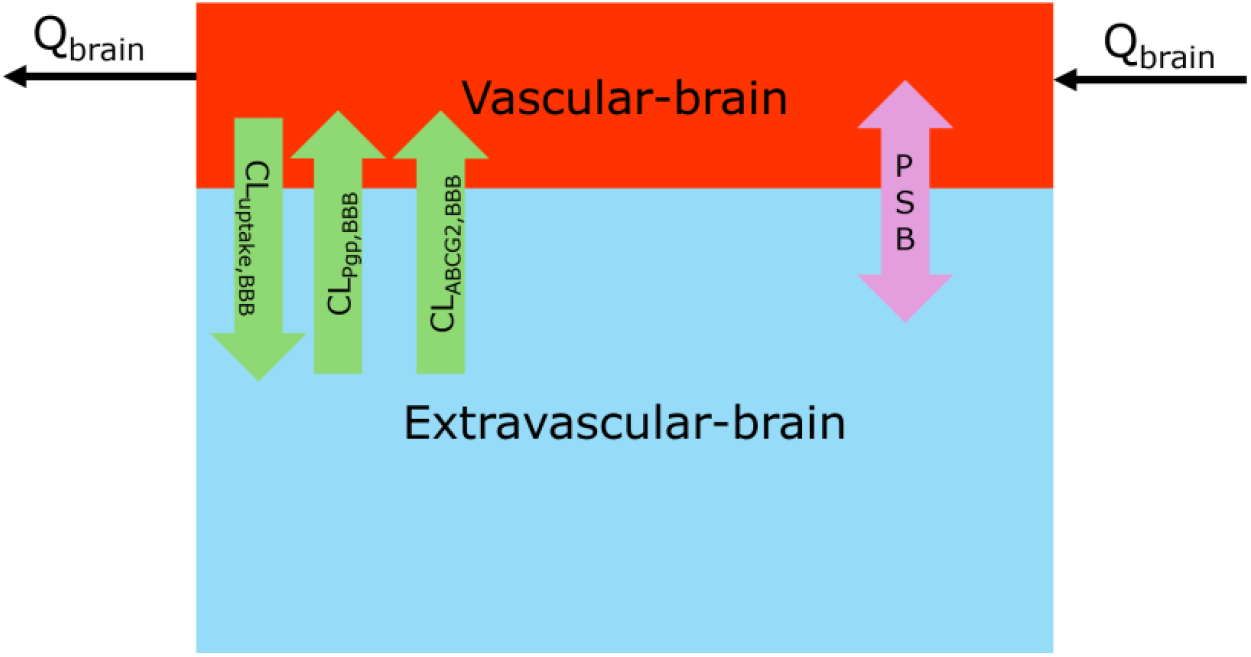
Two-compartment brain model based on structure from Li *et al* (15). There are four membrane transport processes. Three unidirectional BBB pumps – 2 efflux; P-glycoprotein (Pgp) and ABCG2, and 1 uptake – expressed as clearance terms and 1 bidirectional diffusion process (PSB). *Q*_*br*_ is the brain blood or plasma flow rate.

The complete set of sPBPK model equations and parameters are provided in Tables S1 and S2, respectively. The sPBPK AZD1775 model predicted and digitized AZD1775 plasma and brain concentration-time plots are shown in Fig. S2A and S2B, respectively. Agreement between the predicted and digitized plasma and brain areas under the concentration-time curves (AUCs) was high, with the % error = 12.6% for plasma and 3.1% for brain.

### scPBPK Model for AZD1775

The scPBPK model for AZD1775 predicted concentrations were analogous to those produced by the sPBPK model other than those for brain, for example, see Fig. S3 for plasma. The scPBPK model brain structure is shown in Fig. 3. It shows the same membrane transport processes as the sPBPK model, and in addition, the single cell and bulk IC compartments. The ED processes and their inherent variability affect the single cell brain IF and IC concentration profiles (see Figs. 4 and 5, respectively). AZD1775 concentration-time profiles for four clusters - each with a unique weighting function – are shown that illustrates the mean profile for each cluster, as well as bands for the interquartile ranges and 5% - 95% ranges. The brain IF and IC concentrations ranges – based on either the ICR or 5-95% values - vary between clusters with cluster 1 showing the greatest variability of about 4-fold at the maximum concentration values. The NB weighting functions for each cluster were selected randomly between means of 0.5 – 2.5 and dispersion values between 1 (high limit of variability) – 20 (low limit of variability). Through the randomization process cluster 1 attained the greatest variability amongst the 4 clusters. The analogous AZD1775 brain IF and IC bulk concentration-time profiles are shown in Fig. 6, and these profiles more closely track the mean of the single cell profiles although some cluster means are closer than others.

**Fig. 3.**
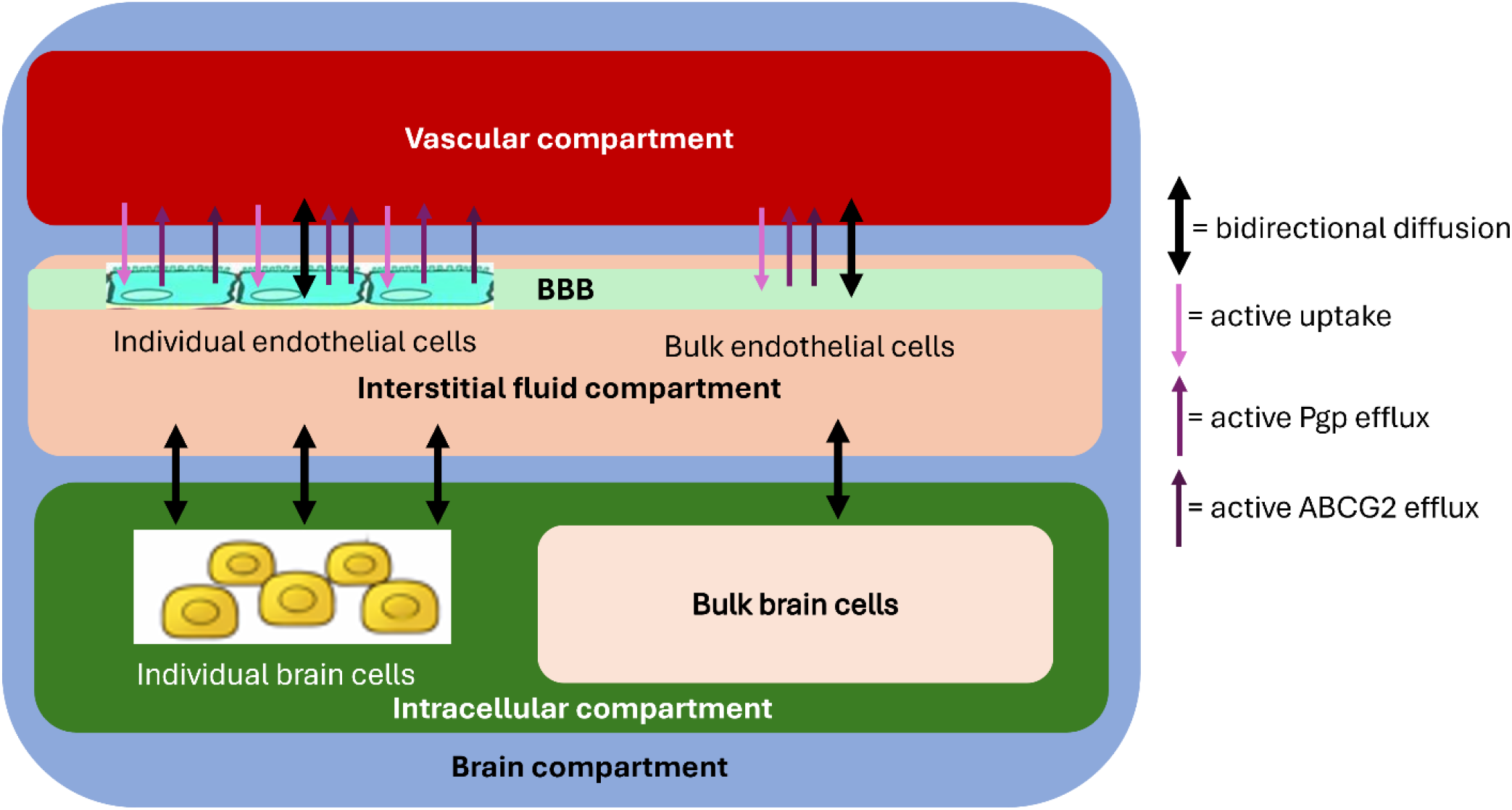
Brain structure for scPBPK model for AZD1775. All membrane transport processes are designated by arrows including the 3 ED processes; active uptake, active Pgp efflux and active ABCG2 efflux. Both the IF and IC compartments have single cell and bulk cell components.

**Fig. 4.**
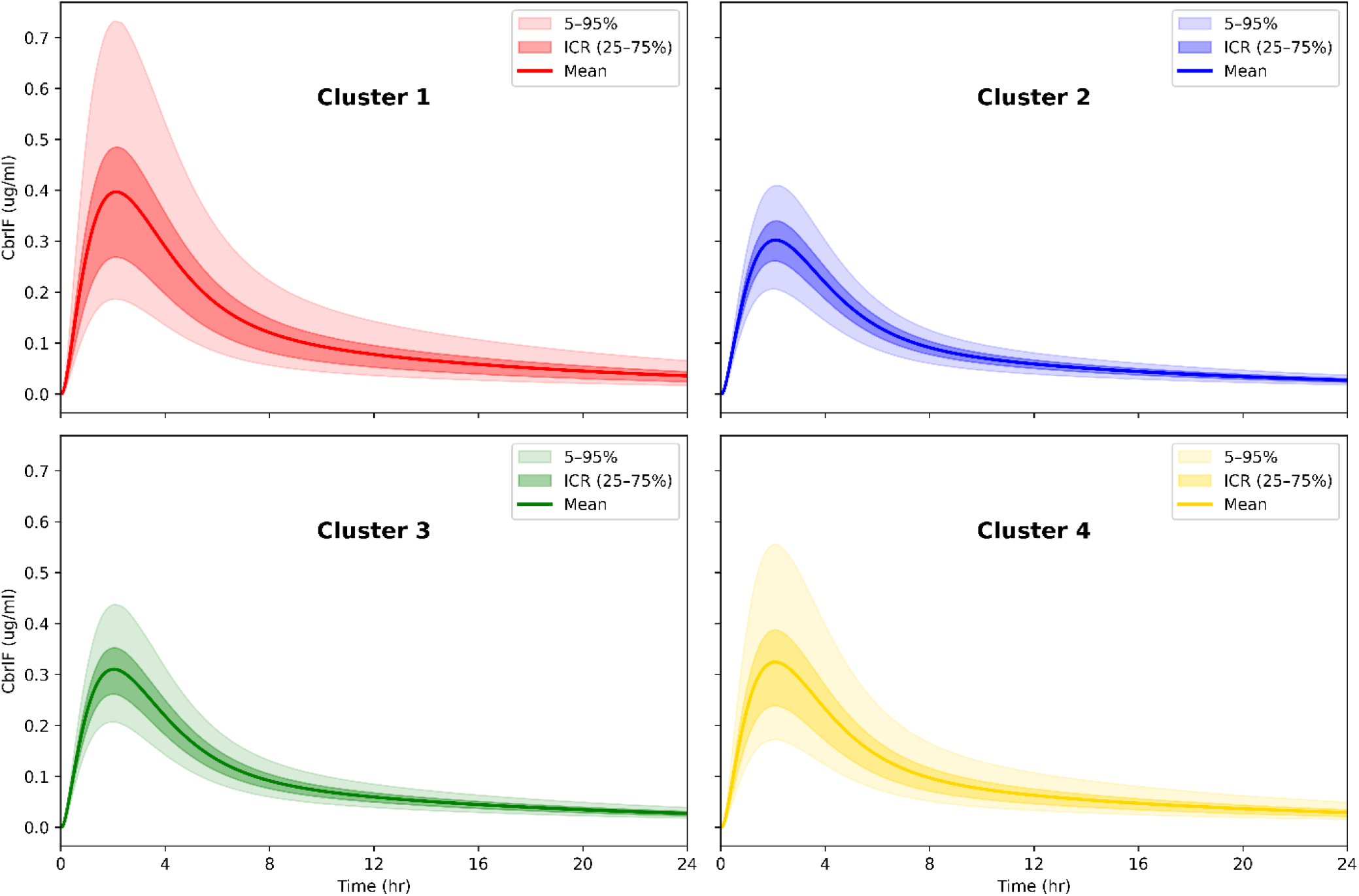
AZD1775 scPBPK model-predicted brain interstitial fluid (IF) single cell concentration-time profiles. The mean concentration, ICR (25-75%) and 5-95% ranges are based on 2500 individual cell profiles per cluster.

**Fig. 5.**
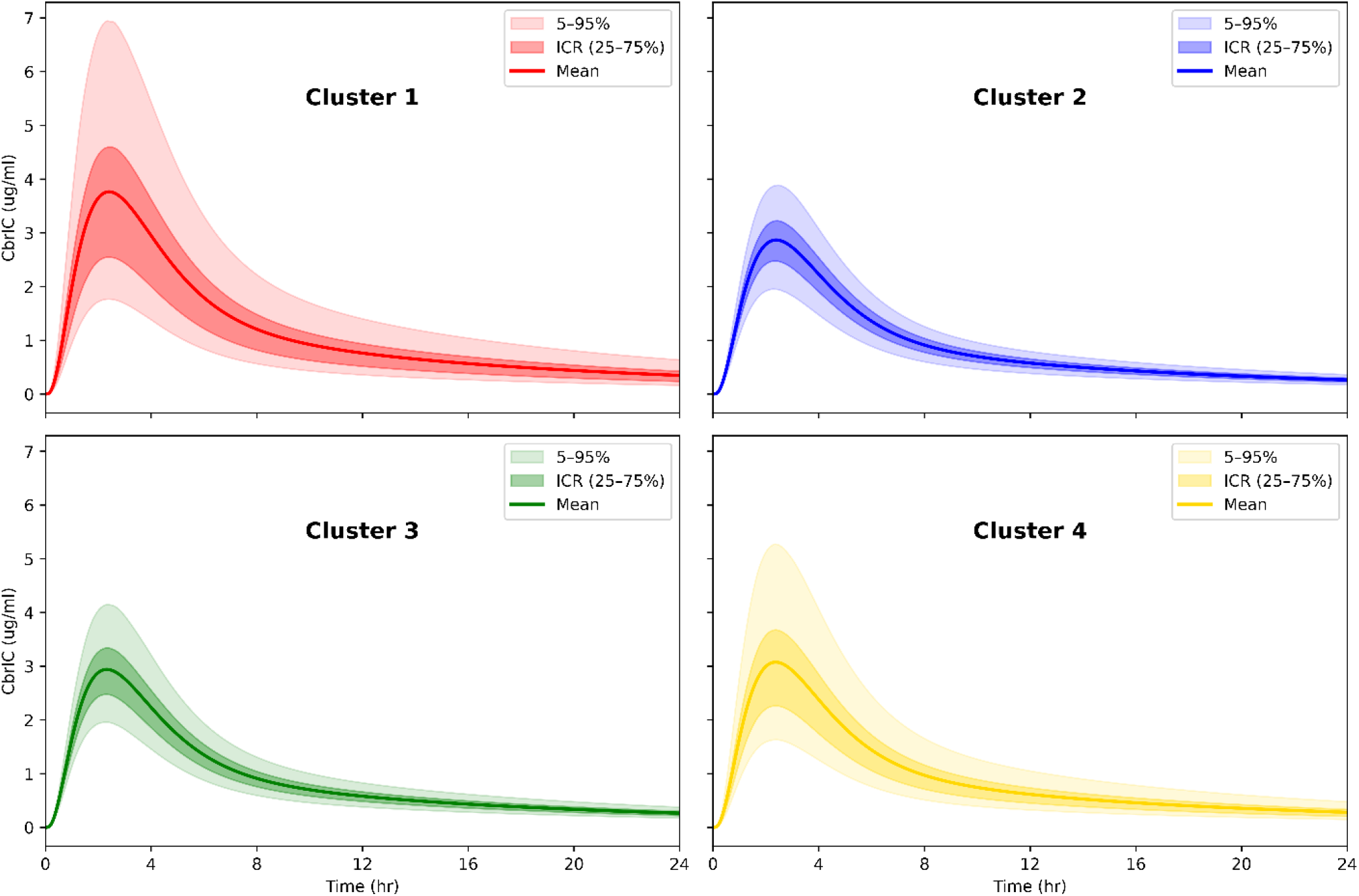
AZD1775 scPBPK model-predicted brain intracellular (IC) single cell concentration-time profiles. The mean concentrations, ICR (25-75%) and 5-95% ranges are based on 2500 individual cell profiles per cluster.

**Fig. 6.**
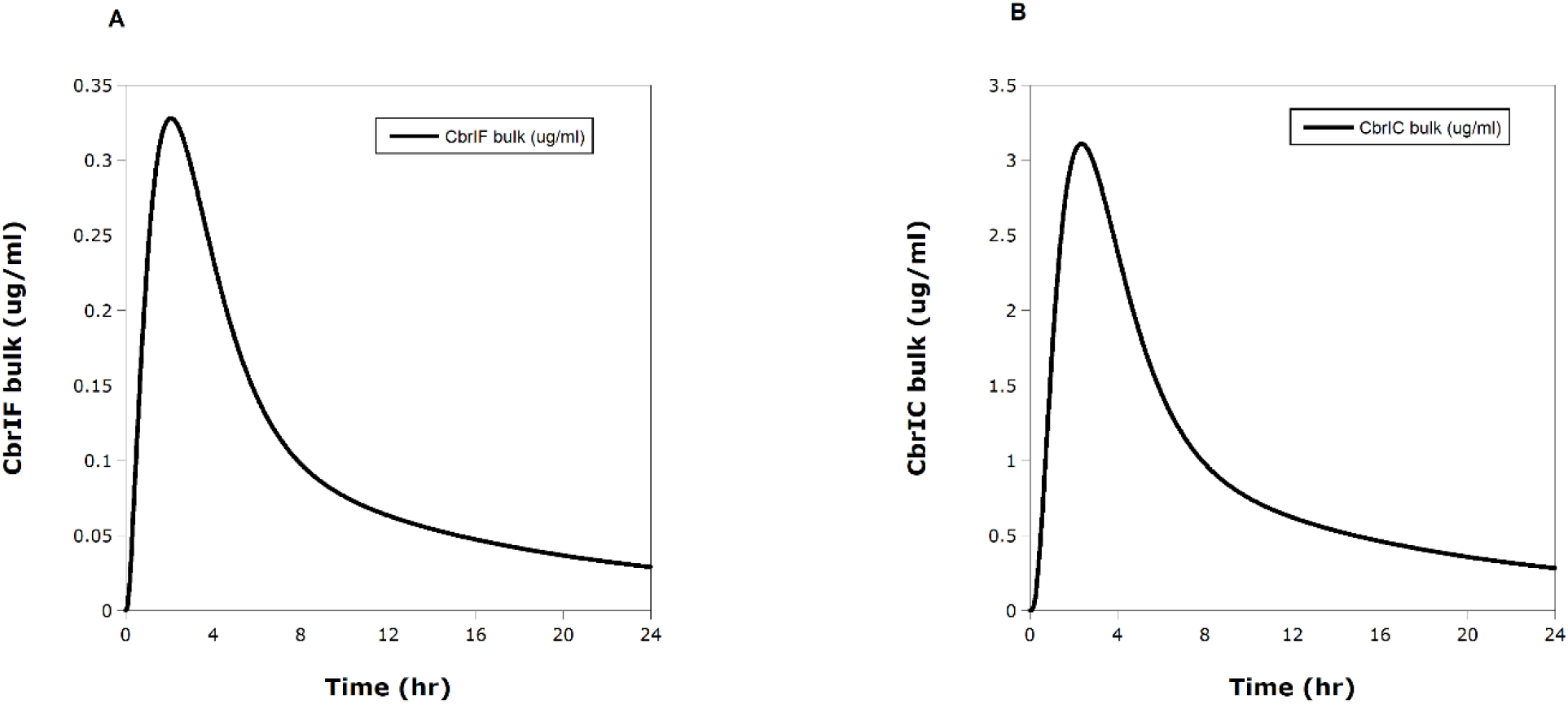
AZD1775 scPBPK model-predicted brain IF (A) and IC (B) bulk AZD1775 concentration-time profiles.

### Minimal PBPK (mPBPK) Model for MDZ

Midazolam (MDZ), a drug categorized as a high clearance drug and used as a probe for CYP3A4 and CYP3A5 metabolism (20, 21) was selected to model as a mPBPK – minimal PBPK – model (22). Minimal PBPK models utilize the phenomenon that organs that behave kinetically the same can be lumped into single generic compartments as noted for midazolam (MDZ) in Fig. 7A that has two lumped compartments, V1 and V2, in addition to a liver compartment that accounts for drug elimination. In our mPBPK model for MDZ we choose to represent the liver compartment as a 3-subcompartment structure to facilitate conversion to a scPBPK model (next section). Drug metabolism of MDZ is primarily due to two key enzymes – CYP3A4 and CYP3A5 – that hydroxylate MDZ to two primary metabolites, 1’-hydroxymidazolam and 4-hydroxymidazolam (20, 23). In many studies (20, 23) MDZ metabolism was considered to be a nonlinear Michaelis-Menton process and was also used in our mPBPK model for MDZ rather than a linear clearance process used by Cao and Jusko (22). Our MDZ mPBPK model-predicted and observed MDZ plasma concentrations (24, see Methods section) are shown in Fig. S4 that indicates accuracy of the model.

**Fig. 7A and 7B.**
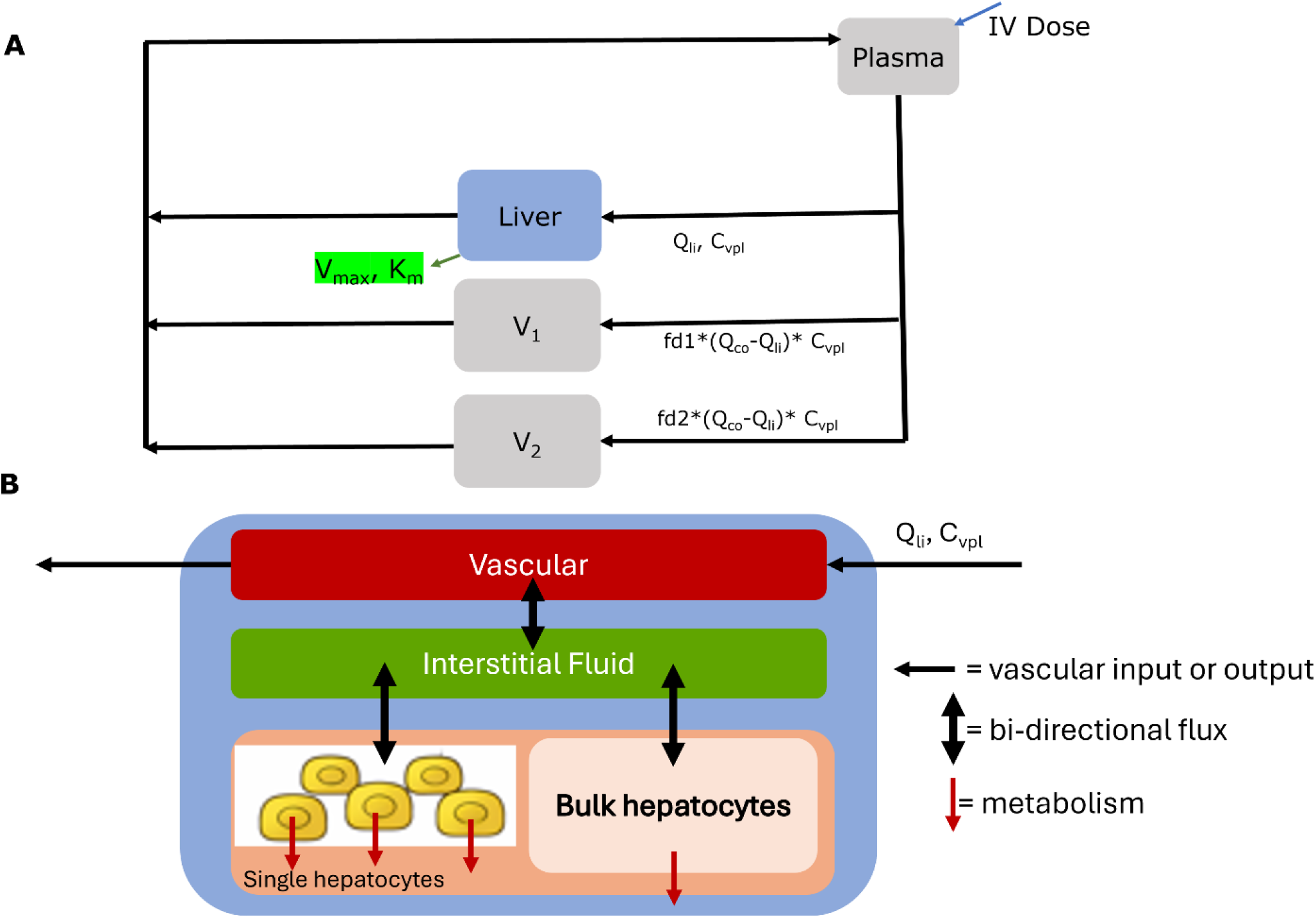
Midazolam (MDZ) mPBPK (A) model and the structure of the liver compartment used in the scPBPK model for MDZ. Q_co_ = cardiac output, Q_li_ = liver plasma flow, all other parameters defined in Table S4.

### scPBPK Model for MDZ

The behavior of the MDZ scPBPK model is unlike that of the AZD1775 scPBPK model in that the liver IC compartment (Fig. 7B) – the compartment with the ED metabolic process – MDZ concentration profiles for all single cells and bulk cells are equal (Figs. 8A), whereas high heterogeneity was observed in the AZD1775 single cell profiles (Figs. 4 and 5). The lack of concentration heterogeneity observed for MDZ liver IC concentrations is due to the high mass transfer coefficients (h_1_ and h_2_, see Table S4) relative to Vmax, a key component of the metabolic ED clearance. In fact, single cell metabolic clearances that depend on Vmax do show cell-to-cell variations (see Fig. 8B). The variation in single cell clearances due to the NB-based weighting functions imparts much greater variation in cluster 1 compared to the other clusters. Nonetheless, the high mass transport of MDZ compared to its metabolic clearance negates the cellular differences in concentrations. When the ratio of h_2_/V_max_ is varied over several orders of magnitude the variances (See Materials and Methods) of MDZ liver IC concentrations exhibit a biphasic relationship (see Fig. 9) over 6 orders of magnitude. The maximum variance occurs at a h2/Vmax ratio of about 0.01. At low ratios there isn’t sufficient drug in the cells where ED metabolic heterogeneity could be operative, and then at high ratios, MDZ transport between the IF and IC compartments is rapid relative to metabolism, which tends to homogenize intracellular concentrations across cells and thereby reduces variance.

**Fig. 8A and 8B.**
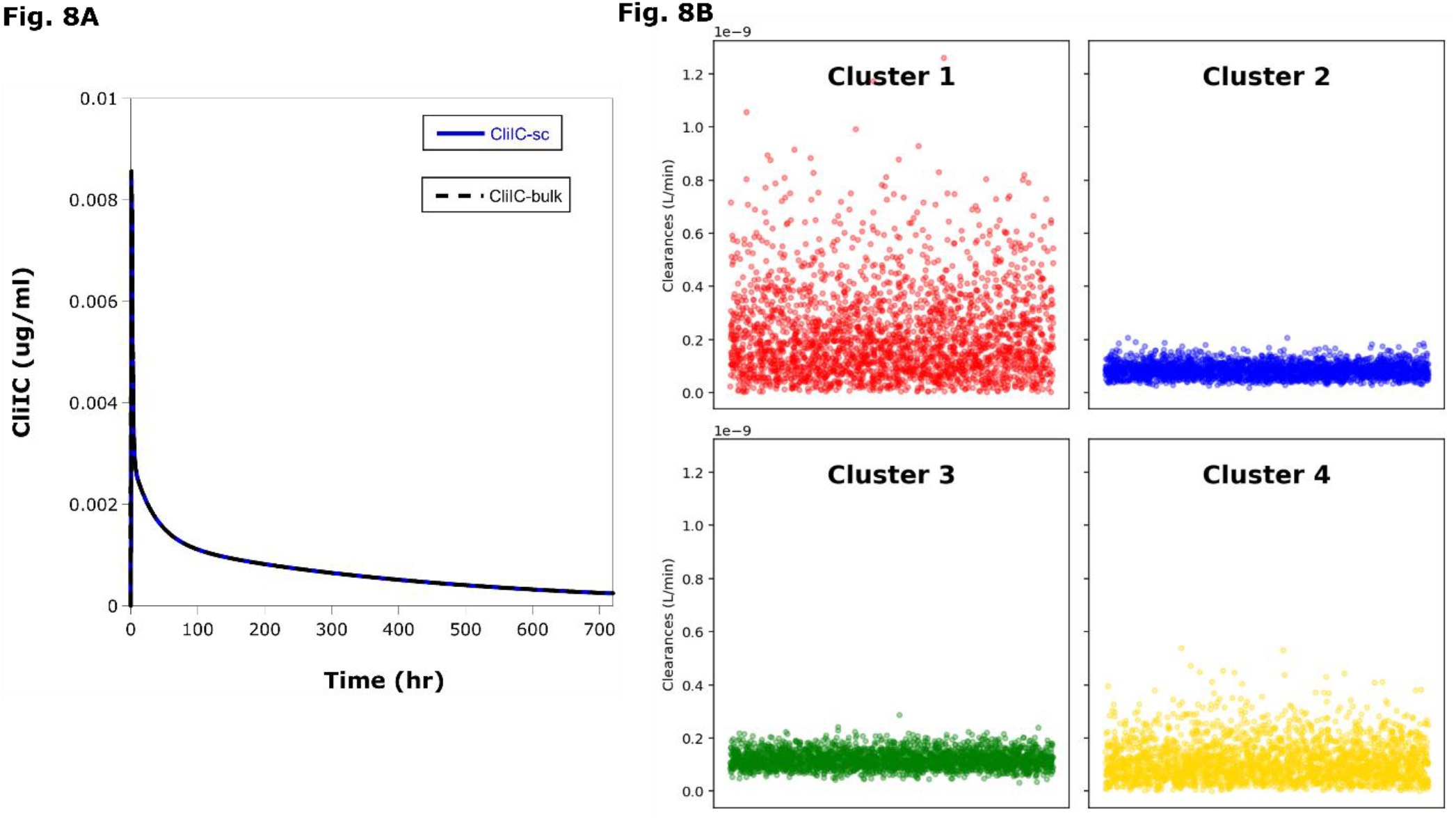
MDZ scPBPK model-predicted liver concentrations (A) and clearances (B). MDZ single cell and bulk liver concentrations profiles are equal (A). Individual single cell clearances vary between each cell and cluster (B). n=2500 for each cluster.

**Fig. 9.**
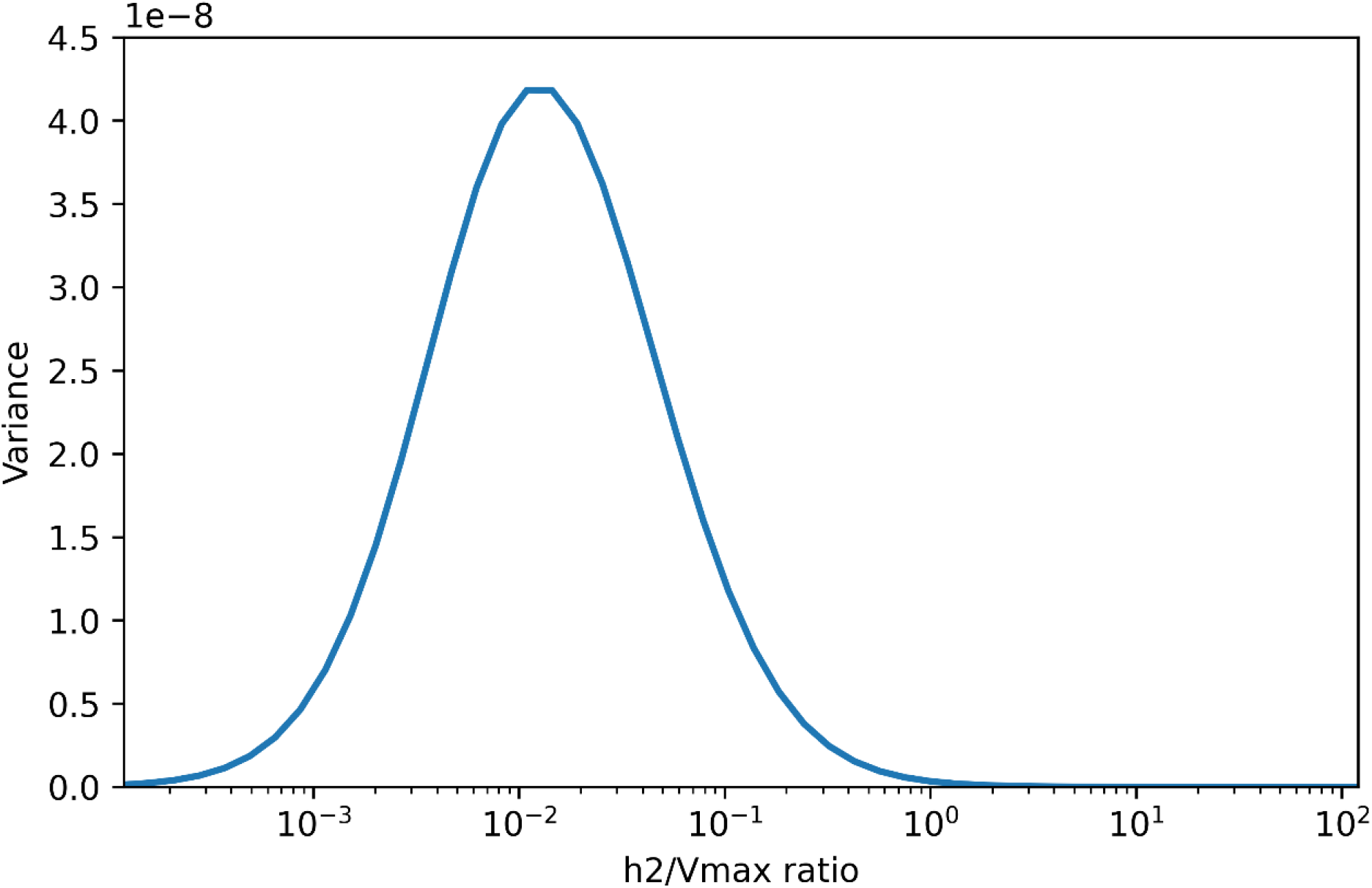
MDZ intracellular liver concentration variance as function of the ratio of h_2_/V_max_. The linear y-axis is based on variances over 6 orders of magnitude that define the tail near 0. The highest variation is at a ratio of 0.011.

## Discussion

scPBPK models offer an opportunity to characterize drug concentration heterogeneity in tissues of interest. This ability can provide important insights as to how individual cells behave and contrast how unique cell types kinetically process drugs. The implications are significant when certain cell types are predicted to have suboptimal drug concentrations indicative of treatment failure. For example, if a greater fraction of cell type 1 or cluster 1 cells are predicted to achieve much greater drug concentrations (*i*.*e*. > EC_50_, for instance) than the fraction of cell type 2 or cluster 2 cells that prevent the intended action on cell type 2 a treatment failure may ensue. Therefore, new strategies could be implemented to overcome the lower concentrations in cell type 2, and such strategies could be designed directly towards cell type 2. Moreover, the scPBPK model used to make such predictions would be able to parse out the mechanism associated with the ED process, such as metabolism or drug transport, and the root cause of lower single cell drug concentrations.

Just as pharmacokinetic (PK) models have evolved to link to pharmacodynamic (PD) models forming PK/PD and PBPK/PD models (25, 26) it is anticipated scPBPK models will also connect to scPD models forming scPBPK/PD models. The field of QSP provides a rich milieu of methods and models to facilitate the development of scPBPK/PD models that could, in addition, to contrasting single cell drug concentrations between clusters can also contrast individual cell responses based on differences in protein dynamics.

Construction of scPBPK models can be built *de novo* or by converting existing sPBPK models as we did here. In either case, the formalism of building sPBPK models apply by grouping system-dependent variables (*i*.*e*. organ blood flows, tissue volumes for species of interest) and drug-dependent variables (*i*.*e*. partition coefficients, mass transfer coefficients, organ clearances). The ED processes must be identified in the particular organ and subcompartment (*i*.*e*. vascular, interstitial fluid, intracellular) and the ODEs delineated. In this regard, choices of a weighting function for each ED process have to be defined either from actual gene or protein expression data or by assuming the nature of the expression distribution as we did here. Fundamental to the scPBPK approach is differentiating single cells from bulk cell dynamics with specific ODEs associated with each. Finally, the parameters associated with the ODEs impacted by the ED processes need to conserve that process over all the cells.

The latter parameter conservation aspect raises key assumptions in building scPBPK models in that the total number of cells (*n*_*tot*_) involved in an ED process must be estimated. Once this value is set, the single cell number (*n*_*sc*_) selected for the simulations – ideally based on the scRNAseq data – will determine the bulk cell number (*n*_*bulk*_). It is also apparent that the scPBPK model predictions are just that and not uniquely grounded in single cell drug concentration measurements. Innovative techniques to measure drug concentrations in single cells are not yet routinely feasible although a custom-made system has reported single cell amounts of the anticancer drug irinotecan (27). It should be appreciated that the whole organ drug concentration measurements that are inherent in building sPBPK models support parameter estimation for scPBPK models as we did here.

The framework for scPBPK models has been presented to provide key insights into cellular PK heterogeneity. Given the current emphasis on multiomic single cell methods and the powerful insights they offer to understanding disease biology, scPBPK models provide a companion tool to unravel single cell pharmacology.

## Materials and Methods

### Fundamental Approach

The framework for the scPBPK approach utilized a simple generic model structure (Fig. 1) to demonstrate the essential features including the ordinary differential equations (ODEs) and parameter definitions.

### Standard PBPK Model for AZD1775

A sPBPK model of AZD1775 was developed prior to converting it to a scPBPK model. The advantage of the AZD1775 investigation was that both plasma and brain AZD1775 concentrations were measured in 12 patients with brain tumors or glioblastoma multiforme (GBM), which permitted us to calibrate our model to actual data (15). Since our ultimate goal was to demonstrate the scPBPK model for AZD1775 we initiated the effort by revising the 2-subcompartment brain structure (Fig. 2) to a 3-subcompartment model (Fig. 3). We relied on the well-known software package PK-Sim^®^ (28, [PK-Sim](https://www.open-systems-pharmacology.org/)), and subsequently, Magnolia ([Magnolia](https://www.magnoliasci.com/)) for our sPBPK model of AZD1775 since it facilitates coding for user-initiated models. PK-Sim^®^ provided organ blood or plasma flow rates, tissue volume and drug-dependent parameters, such as membrane permeability and tissue to plasma partition coefficients based on known physicochemical parameters specific to AZD1775 that supported blood flow-limited tissue models except for brain. The remaining model parameters that included oral bioavailability, kidney and liver elimination, and all the brain-associated parameters were based on those from Li *et al* (15), yet were finalized based on least-squares optimization using their simulated mean AZD1775 plasma and brain concentrations (see Figures 5A and 5C in reference 15) in patients that received a single 400 mg AZD1775 oral dose. The Li *et al* (15) model-predicted mean plasma and brain concentrations agreed with the measured concentrations, and for our model were digitized (Un-Scan-It, version 7, Silk Scientific corporation, [Un-Scan-It](https://www.silkscientific.com/graph-digitizer.htm)) that provided simulated values to provide additional data for fitting. The complete set of sPBPK model equations and parameters are provided in Tables S1 (equations) and S2 (parameters). The base PBPK AZD1775 model predicted and digitized AZD1775 plasma and brain concentration-time plots are shown in Figures S2A and S2B, respectively. Agreement between the predicted and digitized plasma and brain areas under the concentration-time curves (AUCs) was high, with the % error = 12.6% for plasma and 3.1% for brain.

### scPBPK model for AZD1775

The sPBPK model for AZD1775 was converted to a scPBPK model that required new ODEs for the brain compartment (see Fig. 3 and Table S1). The three unidirectional transport processes in the brain compartment – 1 uptake and 2 efflux – were assumed to be ED processes and are reflected in the brain ODEs.

An estimate of the number of brain endothelial cells comprising the BBB was made by using a value of 150 cm^2^/gm (29), the mean surface area of the brain microvasculature, that when multiplied by the brain vascular volume of 30 gm - used in our model – is equal to 4500 cm^2^, which represented the total surface area of the BBB. Next, a single endothelial cell has an estimated surface area of 1.2E-5 cm^2^ (30), and thus, 4500 cm^2^/1.2E-5 cm^2^/cell = 3.7E8 cells.

A negative binomial (NB) distribution was used as the weighting function for each ED process (2 BBB efflux processes and 1 BBB uptake process). Rather than specify a unique mean and theta (dispersion) value for each NB and cluster, a random selection process over means between 0.5 – 2.5 and thetas between 1-20 was used. The dispersion parameter (theta) was sampled on a logarithmic scale between 1 and 20 to avoid bias toward larger theta values, which would otherwise reduce the variance of the NB distribution. In this manner, each weighting function within the four clusters of 2500 cells each has an equal probability of assigning any mean and theta values within these ranges. This randomization process was considered to be less biased compared to preselection of mean and theta values for each cluster.

scPBPK model simulations for both the AZD1775 and MDZ models were completed in Python using Spyder as the IDE.

### Minimal PBPK(mPBPK) Model for MDZ

The mPBPK model for MDZ was based on that used by Cao and Jusko (22) that was calibrated to MDZ plasma concentrations obtained by Heizmann *et al* (24) that administered a single IV bolus dose of 0.15 mg/kg to six healthy volunteers. The model calibration procedure of Cao and Jusko considered alternate model structures and parameters to obtain agreement with the observed MDZ plasma concentrations.

Our mPBPK model for MDZ revised the Cao and Jusko model as follows:

1. a 3-comparment rather than a 1-compartment structure was used for the liver compartment, and
2. a nonlinear Michaelis-Menten formulation was used for intrinsic clearance rather than a linear formulation used by Cao and Jusko (22).

We obtained initial parameter estimates of V_max_ and K_m_ from previously reported in vitro experiments using human liver microsomes (20, 23) and converted these values based on a value of 32 mg of microsomal protein/gm of liver (31) then fit the mPBPK model to the observed plasma MDZ concentrations reported by Heizmann *et al* (24). The mPBPK model for MDZ and the model-predicted and observed MDZ plasma concentrations are shown in Fig. S4. The complete set of mPBPK model equations and parameters are provided in Tables S3 and S4, respectively.

### scPBPK Model for MDZ

The mPBPK MDZ model was converted to a scPBPK model (see Fig. 4A & 4B) that resulted in revisions to the liver ODE system (see Table S4). The scPBPK MDZ model contains one ED process – drug metabolism - in the intracellular liver compartment which required alterations in both the IF and IC ODEs analogous to those presented under the *Fundamental Approach* section. In addition, the number of hepatocytes was estimated to be 1.E8 cells/gm (14), and for the liver intracellular volume of 1.13L or 1130gm then the total number of hepatocytes is estimated as 1.13E11 cells.

The scPBPK model parameters are the same as those used for the mPBPK MDZ model with additional parameters specific to scPBPK MDZ model provided in Table S4.

A negative binomial (NB) distribution was used as the weighting function for the single metabolic clearance ED process. Analogous to the approach taken for the scPBPK model for AZD1775, the weighting function for each cell were randomly selected from a NB distribution with means between 0.5 – 2.3 and thetas between 1-20. The dispersion parameter (theta) was sampled on a logarithmic scale within this range to avoid bias toward larger theta values, which would otherwise result in lower variance in the NB distribution. The weighting values for each cell were then sampled from the corresponding NB distribution. In this manner, each of the four clusters of 2500 cells had an equal probability of receiving any mean and theta values within these ranges.

To investigate the relationships between the IF-IC mass transfer coefficient (*h*_2_) and the maximum metabolic capacity (*V*_*max*_), the scPBPK model for MDZ simulated liver IC concentrations for different values of *h*_2_, incrementally decreasing *h*_2_ so that the *h*_2_/*V*_*max*_ ratio varied over 6 orders of magnitude. The resultant simulated liver IC concentrations were used to obtain the overall mean concentration variance for each *h*_2_/*V*_*max*_ ratio by first calculating the variance in concentration values across cells at each time point and then averaging these variances across all time points.

## Supporting information

"C:\Users\jmgallo\Box\papers\scPBPK\submitted\Supplementary Materials_final.pdf"

## Author Contribution

Conceptualization: JMG, AS

Methodology: JMG, AS

Investigation: JMG, AS

Visualization: JMG, AS Supervision:

JMG Writing—original draft: JMG, AS

## Data and materials availability

The computer code is available on GitHub (https://github.com/anshulsa/scPBPK).

