## Supplementary material for "Introduction to Single-cell Physiologically-Based Pharmacokinetic (scPBPK) Models": "C:\Users\jmgallo\Box\papers\scPBPK\submitted\Supplementary Materials_final.pdf"

**This PDF file includes:**

Figs. S1 to S4

Tables S1 to S4

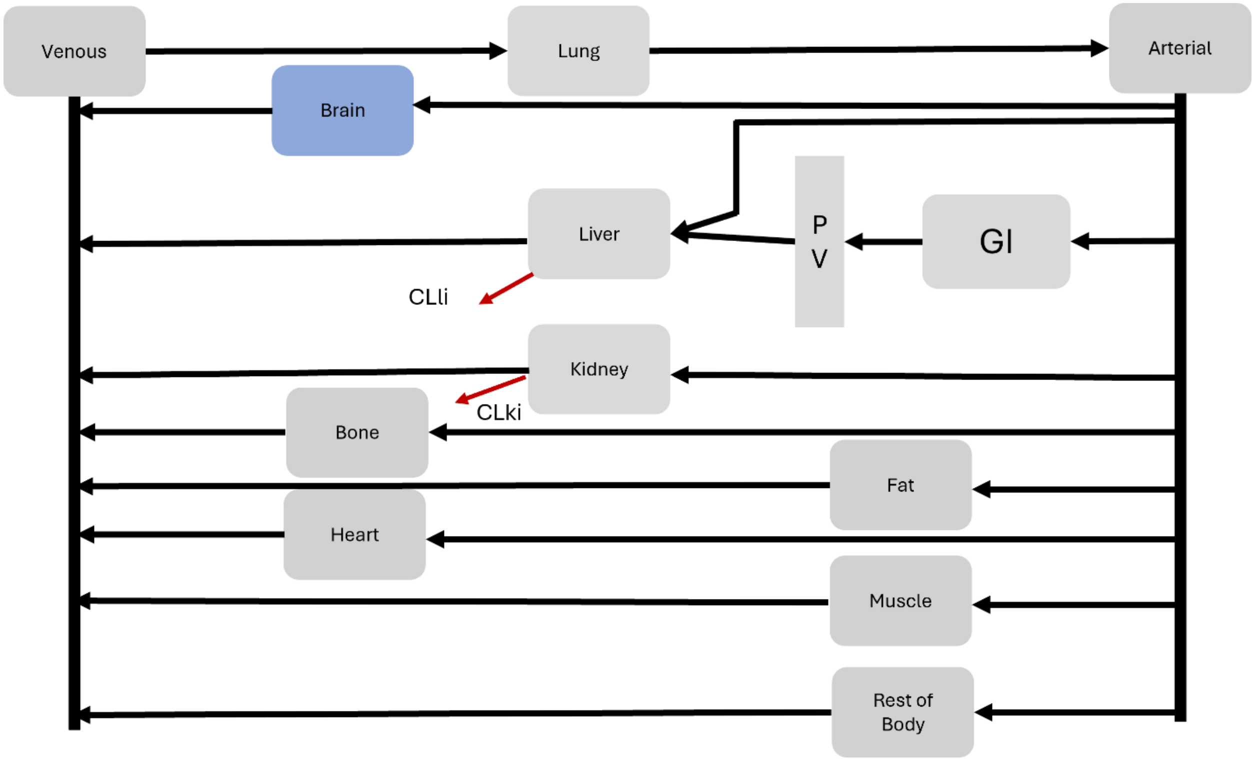

**Fig. S1.** Whole-body sPBPK model for AZD1775. All organs are lumped blood flow-limited compartments except brain, see Fig. 2.

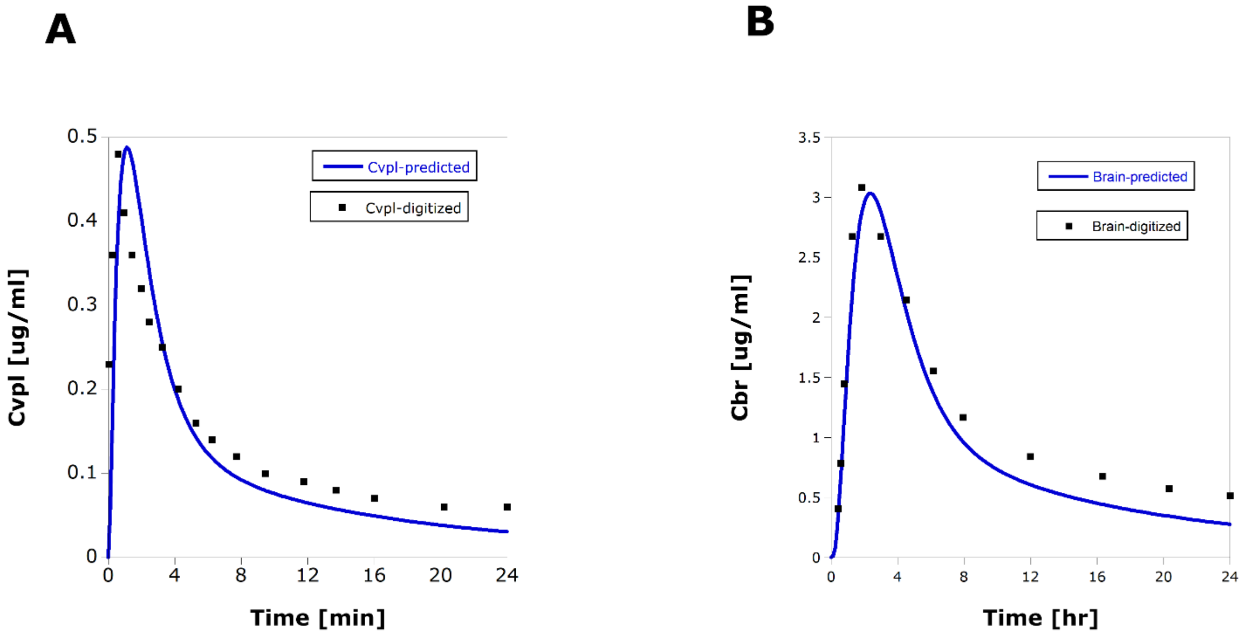

**Fig. S2A and S2B.** sPBPK model-predicted and digitized (observed) AZD1775 plasma (A) and brain (B) concentrations.

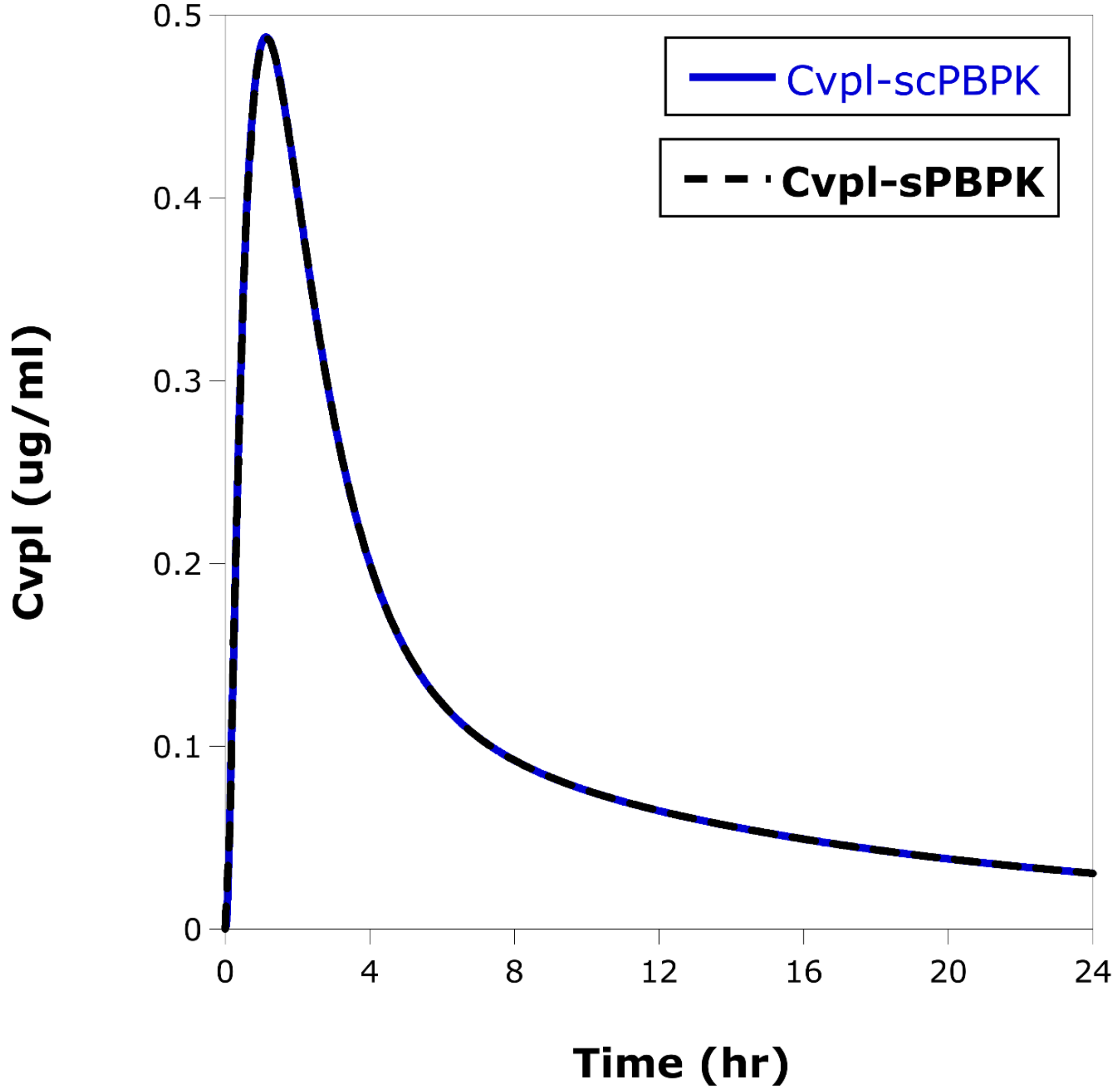

**Fig. S3.** sPBPK and scPBPK model predicted AZD1775 plasma concentrations.

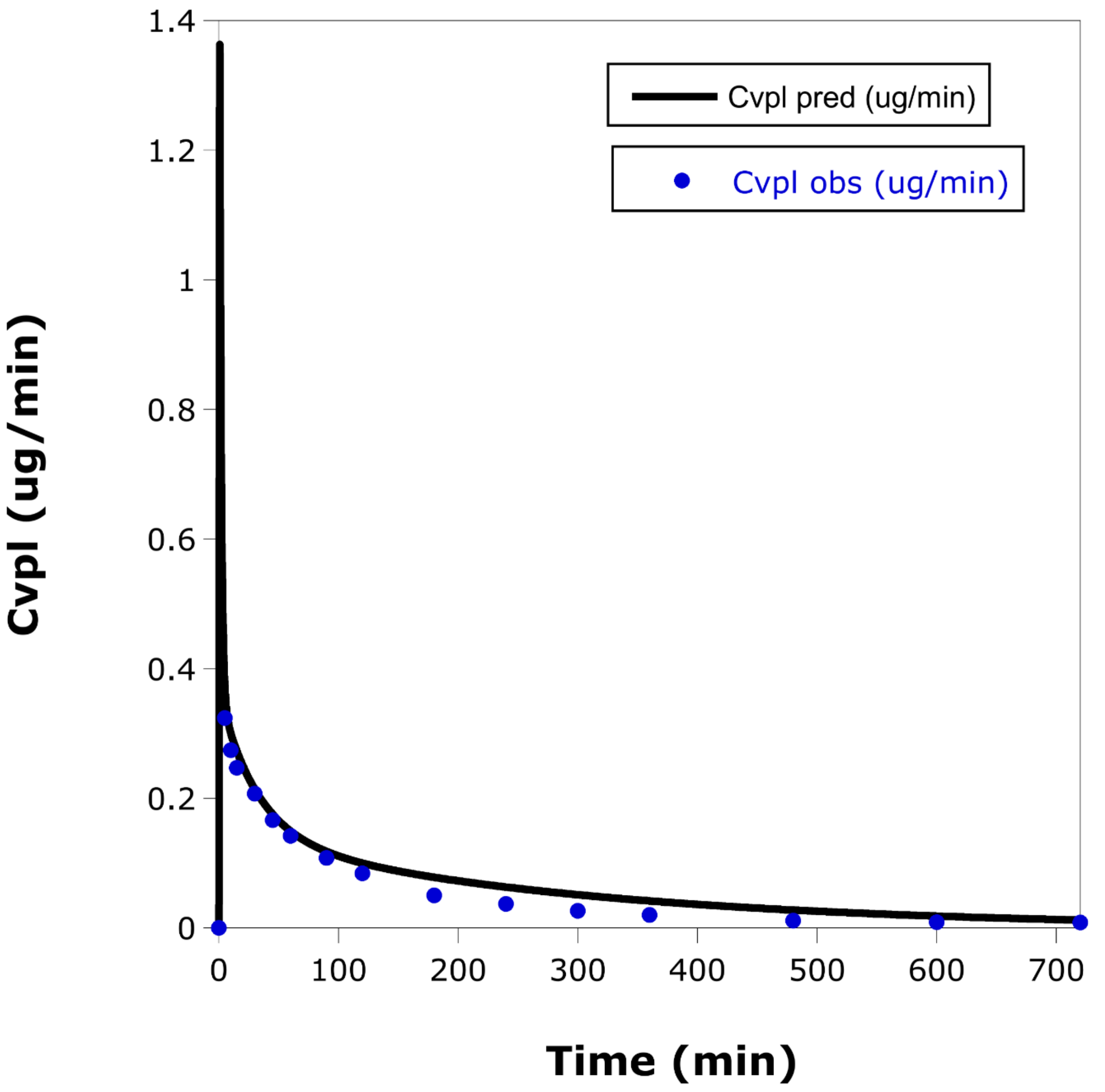

**Fig. S4.** mPBPK model-predicted and observed midazolam plasma concentrations.

**Table S1**. Equations for both the standard PBPK (sPBPK) and scPBPK models for AZD1775. Equations S1-1 thru S1-8 are used for both sPBPK and scPBPK models, Equations S1-9 thru S1-11 are only used for the sPBPK, and Equations S1-12 thru S1-16 are only used for the scPBPK model.

| Equation Number | Compartment | Equation |
|---|---|---|
| S1-1 | Venous plasma(vpl) | $\frac{dA_{vpl}}{dt} = Q_{bo}\frac{C_{bo}}{R_{bo}} + Q_{br}C_{brV} + Q_{fa}\frac{C_{fa}}{R_{fa}} + Q_{he}\frac{C_{he}}{R_{he}} + Q_{ki}\frac{C_{ki}}{R_{ki}} + (Q_{li} + Q_{pv})\frac{C_{li}}{R_{li}} + Q_{mu}\frac{C_{mu}}{R_{mu}} + Q_{rb}\frac{C_{rb}}{R_{rb}} - Q_{co}C_{vpl}$ |
| S1-2 | Lung(lu) | $\frac{dA_{lu}}{dt} = Q_{co}C_{vpl} - Q_{co}\frac{C_{lu}}{R_{lu}}$ |
| S1-3 | Arterial plasma(apl) | $\frac{dA_{apl}}{dt} = Q_{co}\frac{C_{lu}}{R_{lu}} - Q_{co}C_{apl}$ |
| S1-4 | GI(gi) | $\frac{dA_{gi}}{dt} = Q_{gi}C_{apl} - Q_{gi}\frac{C_{gi}}{R_{gi}} + r_{abs}(t)$ |
| S1-5 | Portal vein(pv) | $\frac{dA_{pv}}{dt} = Q_{gi}\frac{C_{gi}}{R_{gi}} - Q_{pv}C_{pv}$ |
| S1-6 | Kidney(ki) | $\frac{dA_{ki}}{dt} = Q_{ki}C_{apl} - Q_{ki}\frac{C_{ki}}{R_{ki}} - CL_{ki,int}\,C_{ki}\left(\frac{f_{up}}{R_{ki}}\right)$ |
| S1-7 | Liver(li) | $\frac{dA_{li}}{dt} = Q_{li}C_{apl} + Q_{pv}C_{pv} - (Q_{li} + Q_{pv})\frac{C_{li}}{R_{li}} - CL_{li,int}\,C_{li}\left(\frac{f_{up}}{R_{li}}\right)$ |
| S1-8 | x=bo,fa,he, mu,rb for bone,fat, heart,muscle or rest of body | $\frac{dA_x}{dt} = Q_xC_{apl} - Q_x\frac{C_x}{R_x}$ |
| S1-9 | Brain vascular (brV) | $\frac{dA_{brV}}{dt} = Q_{br}C_{apl} - Q_{br}C_{brV} - \left[PSB\left(C_{brV} - \frac{C_{brIF}}{R_{br1}}\right) + CL^{upt}f_{up}C_{brV} - (CL^{pgp} + CL^{abcg2})\left(\frac{C_{brIF}}{R_{br1}}\right)\right]$ |
| S1-10 | Brain Interstitial fluid (brIF) | $\frac{dA_{brIF}}{dt} = PSB\left(C_{brV} - \frac{C_{brIF}}{R_{br1}}\right) + CL^{upt}f_{up}C_{brV} - \left[(CL^{pgp} + CL^{abcg2})\left(\frac{C_{brIF}}{R_{br1}}\right)\right] - PSB2\left(C_{brIF} - \frac{C_{brIC}}{R_{br2}}\right)$ |
| S1-11 | Brain intracellular (brIC) | $\frac{dA_{brIC}}{dt} = PSB2\left(C_{brIF} - \frac{C_{brIC}}{R_{br2}}\right)$ |

| | | |
|---|---|---|
| S1-12 | Brain vascular (brV) | $$\frac{dA_{brV}}{dt} = Q_{br}C_{apl} - Q_{br}C_{brV} - \sum_{i=1}^{n_{sc}}\left[PSB_i\left(C_{brV} - \frac{C_{brIF,i}}{R_{br1}}\right) + CL_i^{upt}f_{up}C_{brV} - \left(CL_i^{pgp} + CL_i^{abc}\right)\left(\frac{C_{brIF,i}}{R_{br1}}\right)\right] - \left[PSB_{bulk}\left(C_{brV} - \frac{C_{brIF,bulk}}{R_{br1}}\right) + CL_{bulk}^{upt}f_{up}C_{brV} - \left(CL_{bulk}^{pgp} + CL_{bulk}^{abc}\right)\left(\frac{C_{brIF,bulk}}{R_{br1}}\right)\right]$$ |
| S1-13 | Brain Interstitial fluid (brIF) – single-cell | $$\frac{dA_{brIF,i}}{dt} = PSB_i\left(C_{brV} - \frac{C_{brIF,i}}{R_{br1}}\right) + CL_i^{upt}f_{up}C_{brV} - CL_i^{pgp}\left(\frac{C_{brIF,i}}{R_{br1}}\right) - CL_i^{abcg2}\left(\frac{C_{brIF,i}}{R_{br1}}\right) - PSB2_i\left(C_{brIF,i} - \frac{C_{brIC,i}}{R_{br2}}\right)$$ |
| S1-14 | Brain Interstitial fluid (brIF) – bulk | $$\frac{dA_{brIF,bulk}}{dt} = PSB_{bulk}\left(C_{brV} - \frac{C_{brIF,bulk}}{R_{br1}}\right) + CL_{bulk}^{upt}f_{up}C_{brV} - CL_{bulk}^{pgp}\left(\frac{C_{brIF,bulk}}{R_{br1}}\right) - CL_{bulk}^{abcg2}\left(\frac{C_{brIF,bulk}}{R_{br1}}\right) - PSB2_{bulk}\left(C_{brIF,bulk} - \frac{C_{brIC,bulk}}{R_{br2}}\right)$$ |
| S1-15 | Brain intracellular (brIC) - single-cell | $$\frac{dA_{brIC,i}}{dt} = PSB2_i\left(C_{brIF,i} - \frac{C_{brIC,i}}{R_{br2}}\right)$$ |
| S1-16 | Brain intracellular (brIC) - bulk | $$\frac{dA_{brIC,bulk}}{dt} = PSB2_{bulk}\left(C_{brIF,bulk} - \frac{C_{brIC,bulk}}{R_{br2}}\right)$$ |

*Definitions for sPBPK model:*

$A_z = amount\ in\ compartment\ z;$

$Q_z = plasma\ flow\ rate\ in\ compartment\ z;$

$V_z = volume\ in\ compartment\ z$

$R_z = tissue\ to\ plasma\ partition\ coefficient\ for\ compartment\ z;$

$CL_{ki,int} = intrinsic\ clearance\ for\ kidney;$

$CL_{li,int} = intrinsic\ clearance\ for\ liver;$

$f_{up} = unbound\ fraction\ in\ plasma;$

$k_{abs} = 1st - order\ absorption\ rate\ constant;$

$F = oral\ bioavailability;$

$z = arterial\ plasma\ (apl), brain\ (br), bone(bo), fat(fa), gastrointestinal\ (gi),$

$heart(he), kidney\ (ki), liver\ (li), lung\ (lu), muscle\ (mu), portal\ vein\ (pv), rest\ of\ body\ (rb).$

To define concentrations:

$$C_z = \frac{A_z}{V_z},$$

$$z = arterial\ plasma\ (apl), bone(bo), fat(fa), gastrointestinal\ (gi), heart(he),$$

$$kidney\ (ki), \quad liver\ (li), lung\ (lu), muscle\ (mu), portal\ vein\ (pv), rest\ of\ body\ (rb).$$

$$C_{brV} = \frac{A_{brV}}{V_{brV}}, C_{brIF} = \frac{A_{brIF}}{V_{brIF}}, C_{brIC} = \frac{A_{brIC}}{V_{brIC}}$$

Additional definitions for scPBPK model:

Single-cell parameters, expression independent:

$$PSB_i = \frac{PSB}{n_{tot}}$$

$$PSB2_i = \frac{PSB2}{n_{tot}}$$

All single-cell expression dependent processes are:

$CL_i^z = \frac{CL^z}{n_{tot}} W_i;\ z = upt, pgp, abcg2.$

Bulk parameters:

$$PSB_{bulk} = f_{bulk,cells} PSB$$

$$PSB2_{bulk} = f_{bulk,cells} PSB2$$

$$CL_{bulk}^z = f_{bulk,cells} CL^z;\ z = upt, pgp, abcg2$$

To define concentrations:

$$C_z = \frac{A_z}{V_z},$$

$z = arterial\ plasma\ (apl), bone(bo), fat(fa), gastrointestinal\ (gi), heart(he),$

$kidney\ (ki), liver\ (li), lung\ (lu), muscle\ (mu), portal\ vein\ (pv), rest\ of\ body\ (rb).$

$$C_{brV} = \frac{A_{brV}}{V_{brV}}, C_{brIF,i} = \frac{A_{brIF,i}}{V_{brIF,i}}, C_{brIF,i} = \frac{A_{brIC,i}}{V_{brIC,i}}, C_{brIF,bulk} = \frac{A_{brIF,bulk}}{V_{brIF,bulk}}, C_{brIC,bulk} = \frac{A_{brIC,bulk}}{V_{brIC,bulk}}$$

**Table S2**. Parameter values used in the sPBPK and scPBPK models for AZD1775.

| *Parameter Symbol [units]* | *Definition* | *Values* |
|---|---|---|
| $Q_{bo}$ (L/hr) | Bone plasma flow rate | 9.2 |
| $Q_{br}$ (L/hr) | Brain plasma flow rate | 25.1 |
| $Q_{fa}$ (L/hr) | Fat plasma flow rate | 12.5 |
| $Q_{he}$ (L/hr) | Heart plasma flow rate | 8.6 |
| $Q_{ki}$ (L/hr) | Kidney plasma flow rate | 42.2 |
| $Q_{li}$ (L/hr) | Liver (hepatic artery) plasma flow rate | 15.5 |
| $Q_{lu}$ (L/hr) | Lung plasma flow rate =cardiac output | 200.4 |
| $Q_{mu}$ (L/hr) | Muscle plasma flow rate | 43.6 |
| $Q_{pv}$ (L/hr) | Portal vein flow rate | 36.0 |
| $Q_{gi}$ (L/hr) | Sum of large intestine, pancreas, spleen, small intestine and stomach plasma flowrates | 36.0 |
| Qrb (L/hr) | Rest of body = sum of gonads and skin | 9.7 |
| $Q_{co}$ (L/hr) | Cardiac output | 200.4 |
| $V_{bo}$ (L) | Bone volume | 10.6 |
| $V_{brV}$ (L) | Brain vascular volume | 0.03 |
| $V_{brIF}$ (L) | Brain interstitial fluid volume | 0.01 |
| $V_{brIC}$ (L) | Brain intracellular volume | 1.41 |
| $V_{fa}$ (L) | Fat volume | 18.6 |
| $V_{he}$ (L) | Heart volume | 0.5 |
| $V_{ki}$ (L) | Kidney volume | 0.4 |
| $V_{li}$ (L) | Liver volume | 1.9 |
| $V_{lu}$ (L) | Lung volume | 0.9 |
| $V_{mu}$ (L) | Muscle volume | 30.4 |
| $V_{pv}$ (L) | Portal vein volume | 0.46 |
| $V_{gi}$ (L) | GI volume = sum of large intestine, pancreas, spleen, small intestine and stomach | 1.6 |
| $V_{rb}$ (L) | Rest of body volume = gonads and skin | 3.5 |
| $V_{apl}$ (L) | Volume of arterial plasma | 0.22 |
| $V_{vpl}$ (L) | Volume of venous plasma | 0.5 |
| $R_{bo}$ | Bone partition coefficient | 129 |
| $R_{br1}$ | Brain partition coefficient between vascular and interstitial compartments | 0.49 |
| $R_{br2}$ | Brain partition coefficient between interstitial and intracellular compartments | 9.6 |
| $R_{fa}$ | Fat partition coefficient | 376 |
| $R_{he}$ | Heart partition coefficient | 49 |
| $R_{ki}$ | Kidney partition coefficient | 26 |

| | | |
|---|---|---|
| $R_{li}$ | Liver partition coefficient | 4.3 |
| $R_{lu}$ | Lung partition coefficient | 7 |
| $R_{mu}$ | Muscle partition coefficient | 8 |
| $R_{gi}$ | GI partition coefficient | 28* |
| $R_{rb}$ | Rest of body partition coefficient | 33[#] |
| $CL_{Pgp,BBB}$ (L/hr) | Efflux clearance due to P-glycoprotein at the BBB | 52.2 |
| $CL_{ABCG2,BBB}$ (L/hr) | Efflux clearance due to ABCG2 at the BBB | 57.6 |
| $CL_{uptake,BBB}$ (L/hr) | Uptake clearance at BBB | 7.4 |
| PSB (L/hr) | Bidirectional transport at BBB | 12.6 |
| PSB2 (L/hr) | Bidirectional transport at IF-IC membrane | 54.2 |
| $K_{abs}$ (hr-1) | Absorption rate constant | 2.1 |
| F | Oral bioavailability | 0.7 |
| $CL_{liInt}$ (L/hr) | Liver intrinsic clearance | 250 |
| $CL_{kiInt}$ (L/hr) | Kidney intrinsic clearance | 5.5 |
| $f_{up}$ | Unbound fraction in plasma | 0.2 |
| | *Parameters Specific to the scPBPK AZD1775 Model* | |
| $n_{tot}$ | Total number of brain endothelial cells forming the BBB | 4.5E8 |
| $n_{sc}$ | Total number of single cells | 10000 |
| $n_{bulk}$ | Total number of bulk cells | 4.49E8 |
| $PSB_i$(L/hr) | Bidirectional transport at BBB for single cells | 2.8E-8 |
| $PSB2_i$(L/hr) | Bidirectional transport at IF-IC membrane for single cells | 1.2E-7 |
| $PSB_{bulk}$(L/hr) | Bidirectional transport at BBB for bulk cells | 12.5 |
| $PSB2_{bulk}$(L/hr) | Bidirectional transport at IF-IC membrane for bulk cells | 54. |
| $CL_i^{upt}$(L/hr) | Uptake clearance at BBB for single cells | 1.64E-8*$W_i$ |
| $CL_i^{pgp}$(L/hr) | Efflux clearance due to P-glycoprotein at the BBB for single cells | 1.16E-7*$W_i$ |
| $CL_i^{abcg2}$(L/hr) | Efflux clearance due to ABCG2 at the BBB for single cells | 1.26E-7*$W_i$ |
| $CL_{bulk}^{upt}$(L/hr) | Uptake clearance at BBB for bulk cells | 7.37 |
| $CL_{bulk}^{pgp}$(L/hr) | Efflux clearance due to P-glycoprotein at the BBB for bulk cells | 52. |
| $CL_{bulk}^{abcg2}$(L/hr) | Efflux clearance due to ABCG2 at the BBB for bulk cells | 57.4 |

Definitions for scPBPK model:

Single-cell parameters, expression independent:

$$PSB_i = \frac{PSB}{n_{tot}}$$

$$PSB2_i = \frac{PSB2}{n_{tot}}$$

All single-cell expression dependent processes are:

$CL_i^z = \frac{CL^z}{n_{tot}} W_i;\ z = upt, pgp, abcg2.$

Bulk parameters:

$$PSB_{bulk} = f_{bulk,cells} PSB$$

$$PSB2_{bulk} = f_{bulk,cells} PSB2$$

$$CL_{bulk}^z = f_{bulk,cells} CL^z;\ z = upt, pgp, abcg2$$

**Table S3**. Equations for both the standard PBPK (sPBPK) and scPBPK models for midazolam (MDZ). Equations S3-1 thru S3-4 are used for both sPBPK and scPBPK models, Equations S3-5 and S3-6 are only for the sPBPK, and Equations S3-7 thru S3-9 are only used for the scPBPK model.

| Equation Number | Compartment | Equation |
|---|---|---|
| S3-1 | Venous plasma(vpl) | $\frac{dA_{vpl}}{dt} = Q_{li}C_{liV} + f_{d1}(Q_{co} - Q_{li})\frac{C_1}{R_1} + f_{d2}(Q_{co} - Q_{li})\frac{C_2}{R_2} + Dose_{IV} -(Q_{li}C_{vpl} + f_{d1}(Q_{co} - Q_{li})C_{vpl} + f_{d2}(Q_{co} - Q_{li})C_{vpl}$ |
| S3-2 | Lumped 1(1) | $\frac{dA_1}{dt} = f_{d1}(Q_{co} - Q_{li})C_{vpl} - f_{d1}(Q_{co} - Q_{li})\frac{C_1}{R_1}$ |
| S3-3 | Lumped 2(2) | $\frac{dA_2}{dt} = f_{d2}(Q_{co} - Q_{li})C_{vpl} - f_{d2}(Q_{co} - Q_{li})\frac{C_2}{R_2}$ |
| S3-4 | Liver vascular (liV) | $\frac{dA_{liV}}{dt} = Q_{li}C_{vpl} - Q_{li}C_{liV} - h_1\left(C_{liV} - \frac{C_{liIF}}{R_{li1}}\right)$ |
| S3-5 | Liver interstitial fluid (liIF) | $\frac{dA_{liIF}}{dt} = h_1\left(C_{liV} - \frac{C_{liIF}}{R_{li1}}\right) - h_2\left(C_{liIF} - \frac{C_{liIC}}{R_{li2}}\right)$ |
| S3-6 | Liver intracellular (liIC) | $\frac{dA_{liIC}}{dt} = h_2\left(C_{liIF} - \frac{C_{liIC}}{R_{li2}}\right) - V_{max}\frac{C_{liIC}}{K_m\left(\frac{R_{li2}}{f_{up}}\right) + C_{liIC}}$ |
| S3-7 | Liver interstitial fluid – single-cell model (liIF) | $\frac{dA_{liIF}}{dt} = h_1\left(C_{liV} - \frac{C_{liIF}}{R_{li1}}\right) - \sum_{i=1}^{n_{sc}} h_{2,sc}\left(C_{liIF} - \frac{C_{liIC,i}}{R_{li2}}\right) - h_{2,bulk}\left(C_{liIF} - \frac{C_{liIC,bulk}}{R_{li2}}\right)$ |
| S3-8 | Liver intracellular- single-cell (liIC,i) | $\frac{dA_{liIC,i}}{dt} = h_{2,sc}\left(C_{liIF} - \frac{C_{liIC,i}}{R_{li2}}\right) - V_{max,i}\frac{C_{liIC,i}}{K_m\left(\frac{R_{li2}}{f_{up}}\right) + C_{liIC,i}}$ |
| S3-9 | Liver intracellular - bulk (liIC,bulk) | $\frac{dA_{liIC,bulk}}{dt} = h_{2,bulk}\left(C_{liIF} - \frac{C_{liIC,bulk}}{R_{li2}}\right) - V_{max,bulk}\frac{C_{liIC,bulk}}{K_m\left(\frac{R_{li2}}{f_{up}}\right) + C_{liIC,bulk}}$ |

*Definitions for sPBPK model:*

$A_z = amount\ in\ compartment\ z;$

$Q_z = plasma\ flow\ rate\ in\ compartment\ z;$

$V_z = volume\ in\ compartment\ z$

$R_z = tissue\ to\ plasma\ partition\ coefficient\ for\ compartment\ z;$

$V_{max} = maximum\ metabolic\ rate;$

$K_m = michaelis\ constant\ for\ metabolism;$

$f_{up} = unbound\ fraction\ in\ plasma;$

$$z = \ venous\ plasma(vpl), lumped\ compartment\ 1\ (1), lumped\ compartment\ 2\ (2),$$

$$liver\ (li).$$

To define concentrations:

$$C_z = \frac{A_z}{V_z},$$

$$z = \text{venous plasma(vpl), lumped compartment 1 (1), lumped compartment 2 (2),}$$

$$\text{liver vascular (liV), liver interstitial fluid (liIF ), liver intracellular (liIC).}$$

$$C_{liV} = \frac{A_{liV}}{V_{liV}},\ C_{liIF} = \frac{A_{liIF}}{V_{liIF}},\ C_{liIC} = \frac{A_{liIC}}{V_{liIC}},\ \ C_{v1} = \frac{A_{v1}}{V_1},\ C_{v2} = \frac{A_{v2}}{V_2},$$
$$C_{vpl} = \frac{A_{vpl}}{V_{vpl}}$$

Other definitions:

$$IVDose_{input} = Dose * IV_{smooth}$$

$$IV_{smooth} = 30\frac{t^2}{D^3}\left(1 - \frac{t}{D}\right)^2 Indicator$$

and $Indicator = 1, for\ 0\ \leq t \leq D, else\ 0$

Additional definitions for scPBPK model:

$$h_{2,bulk} = f_{bulk,cells} h_2$$

$$h_{2,sc} = \frac{h_2}{n_{tot}}$$

$$V_{max,bulk} = f_{bulk,cells} V_{max}$$

and

$$f_{bulk,cells} = \frac{n_{bulk}}{n_{tot}}$$

To define concentrations:

$$C_{liV} = \frac{A_{liV}}{V_{liV}},\ C_{liIF} = \frac{A_{liIF}}{V_{liIF}},\ C_{liIC,i} = \frac{A_{liIC,i}}{V_{liIC,i}},\ C_{liIC,bulk} = \frac{A_{liIC,bulk}}{V_{liIC,bulk}},\ C_{v1} = \frac{A_{v1}}{V_1},\ C_{v2} = \frac{A_{v2}}{V_2},$$

$$C_{vpl} = \frac{A_{vpl}}{V_{vpl}}$$

**Table S4**. Parameter values used in the sPBPK and scPBPK models for Midazolam.

| *Parameter Symbol [units]* | *Definition* | *Values* |
|---|---|---|
| $Q_{li}$ (L/min) | Liver plasma flowrate | 1.45 |
| $Q_{co}$ (L/min) | Cardiac output | 5.6 |
| fd1 | Fractional plasma flowrate to compartment 1 | 0.692 |
| fd2 | Fractional plasma flowrate to compartment 2 | 0.0842 |
| $V_{li}$ (L) | Volume of liver compartment | 1.69 |
| $V_{liV}$ (L) | Volume of liver vascular compartment | 0.29 |
| $V_{liIF}$ (L) | Volume of liver interstitial compartment | 0.27 |
| $V_{liIC}$ (L) | Volume of liver intracellular compartment | 1.13 |
| $V_1$ (L) | Volume of compartment 1 | 24.3 |
| $V_2$ (L) | Volume of compartment 2 | 38.4 |
| $V_{vpl}$ (L) | Volume of venous plasma | 5.2 |
| $h_1$ [L/min] | Mass transfer coefficient between vascular and interstitial fluid compartments | 27416. |
| $h_2$ [L/min] | Mass transfer coefficient between interstitial fluid and intracellular compartments | 14606 |
| $R_{li1}$ (dimensionless) | Liver partition coefficient between vascular and interstitial fluid compartments | 0.39 |
| $R_{li2}$ (dimensionless) | Liver partition coefficient between interstitial fluid and intracellular compartments | 0.022 |
| $R_1$ (dimensionless) | Partition coefficient for compartment 1 | 0.655 |
| $R_2$ (dimensionless) | Partition coefficient for compartment 1 | 0.655 |
| $V_{max}$ (mg/min) | Maximum metabolic rate | 12.16 |
| $K_m$ (mg/L) | Michaelis constant | 1.08 |
| $f_{up}$ | Unbound fraction in plasma | 0.03 |
| *Parameters Specific to the scPBPK MDZ Model* | | |
| $n_{tot}$ | Total number of liver hepatocytes | 1.130E11 |
| $n_{sc}$ | Total number of single cells | 10000 |
| $n_{bulk}$ | Total number of bulk cells | 1.129E11 |
| $h_{2,sc}$ [L/min] | Mass transfer coefficient between interstitial fluid and intracellular compartments for single cells | 1.29E-7 |
| $h_{2,bulk}$ [L/min] | Mass transfer coefficient between interstitial fluid and intracellular compartments for bulk cells | 14591 |
| $V_{max,i}$ [mg/min] | Maximum metabolic rate for single cell i | 1.07E-10 |
| $V_{max,bulk}$ [mg/min] | Maximum metabolic rate for bulk cells | 12.15 |

* Equations for parameter:

$$h_{2,sc} = \frac{h_2}{n_{tot}}$$

$$h_{2,bulk} = f_{bulk,cells} h_2$$

$$V_{max,i} = \frac{V_{max}}{n_{tot}} W_i$$

$$V_{max,bulk} = f_{bulk,cells} V_{max}$$
